# Learning Free Energy Pathways through Reinforcement Learning of Adaptive Steered Molecular Dynamics

**DOI:** 10.1101/2022.10.04.510845

**Authors:** Nicholas Ho, John Kevin Cava, John Vant, Ankita Shukla, Jake Miratsky, Pavan Turaga, Ross Maciejewski, Abhishek Singharoy

## Abstract

In this paper, we develop a formulation to utilize reinforcement learning and sampling-based robotics planning to derive low free energy transition pathways between two known states. Our formulation uses Jarzynski’s equality and the stiff-spring approximation to obtain point estimates of energy, and construct an informed path search with atomistic resolution. At the core of this framework, is our first ever attempt we use a policy driven adaptive steered molecular dynamics (SMD) to control our molecular dynamics simulations. We show that both the reinforcement learning and robotics planning realization of the RL-guided framework can solve for pathways on toy analytical surfaces and alanine dipeptide.

## 1 Introduction

Understanding both the structure and dynamics of proteins is crucial for solving problems in molecular biology. Addressing this need, Molecular dynamics, or MD, is a simulation method widely applied to biomolecular systems. However, dynamics often prove to be incredibly difficult to model due to both the high dimensionality of the systems, and the timescale in which most biological processes take place. Steered molecular dynamics (SMD) therefore applies external steering forces in the target direction to accelerate processes that otherwise, due to energy barriers, are too slow. Since the inception of these simulations, a number of brute-force [13, 21] and adaptive [28] variations of this methodology are now available.

Borrowed from the disruptive area of enhanced sampling simulations [26], a first step in any SMD protocol is to determine a reasonable set of collective variables (CVs) that captures the problem of interest. Essentially, these CVs are are a low dimensional representation for the dynamics of the molecule, used to approximate its key degrees of freedom. The selection, screening and learning of CVs mark an active area of important sampling methodologies [6] that intrinsically benefits the SMD formulation. The resulting SMD trajectory is a reflection of conformational transitions on an underlying energy landscape. Yet, this energy information is not apparent in the SMD computations. The non-equilibrium work accumulated of a trajectory has been used as a metric for determining the the quality of the transition pathway [19]. This quantity (defined below in equation 4) is noisy and, by itself has little statistical significance. The famous Jarzynski’s equality [17] connects this nonequilibrium work function to equilibrium averages from Boltzmann distributions, namely free-energy estimates. Despite this milestone and several MD applications [11], free energy, by definition remains a state-function. So Jarzynski’s estimates offers end-point energy differences from a trajectory, and has minimal bearing on the quality of the pathway. So an ensemble of SMD trajectories with Jarzynski’s equality at best offers free energy differences, with no idea about the pathway, creating a major bottleneck in the applicability of external forces in enhanced sampling.

Significant effort has been dedicated to simulating the likely pathways for conformational transition. Prior methods include anisotropic network models [7], steepest decent dynamics [10], and string simulations with swarm of trajectories[20]. We refer to low-energy transition pathways and biologically relevant pathways interchangeably, as these have been analytically found to be the one with least resistance to transition. Previous works have also using generalized ensemble methods sample the entire phase space before determining the low energy pathways [3]. However, with reinforcement learning (RL), we are interested in finding these pathways without the need of learning the entire free energy landscape.

In this paper, we present a novel methodology that uses Jarzynski’s equality and SMD to derive probable transition pathways between two known states. We demonstrate the application of this framework through a policy gradient RL algorithm that systematically explores the free energy landscape to construct such pathways. We then also apply the framework to a robotics planning algorithm RRT^*^ in order to do a comparative analysis between the two algorithms in the context of this problem. As proof of concept, we apply this framework on two toy systems i) a 2-dimensional analytical free energy measurement known with varying degrees of uncertainty, and ii) alanine dipeptide, with 22×3 degrees of freedom, wherein brute-force SMD has always failed to determine plausible pathways.

## 2 Background and Related Work

### 2.1. Jarzynski’s equality, Steered Molecular Dynamics and Stiff Spring Approximation

We will use a form of adaptive SMD to drive the biomolecule from an initial equilibrium state *A* to a final equilibrium state *B*. The bias potential *h*_*λ*_ that drives the MD simulation is based on the steering parameter *λ*, which is determined by the action *a*_*t*_ chosen by the policy at timestep *t. h*_*λ*_ is expressed as

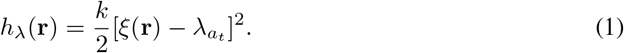

Here *k* is the spring constant used for steering, *ξ* represent one or more CVs that are functions of **r** atomic positions. We express the total Hamiltonian as

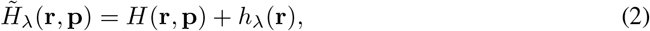

where *H* is the Hamiltonian for the molecular force field and **p** is the momenta. Introducing Jarzynski’s equality, we write

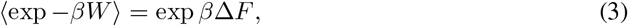

where *β* = 1*/k*_*b*_*T*, T is the temperature and *k*_*b*_ is the Boltzmann constant. The work *W* done by the system along a trajectory is

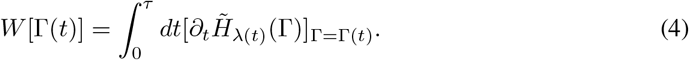

Here Γ(*t*) is the trajectory between state A and state B that evolves *λ*. Approximating the exponential average exp *βW* can be difficult due to small work values that arise only rarely. In order for an accurate measurement of the free energy difference, there needs to be sufficient sampling of rare trajectories that result in small work values [21]. So instead of exponential average, Biophysicists have employed the cumulant expansion for free energy which follows

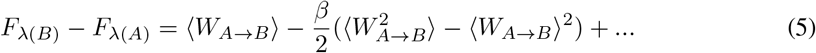

It has been shown that if we use SMD to pull the system along a particular collective variable of interest with a large enough spring constant *k*, then the distribution of work will be Gaussian [21]. This allows us to truncate the approximation of the culmulant expansion to the first two terms for an accurate approximation of the free energy difference. Intuitively, one would choose the spring constant *k* to be large enough such that fluctuations along the trajectory are minimized. In summary, we use this stiff-spring approximation alongside Jarzynski’s equality to obtain free energy estimates. These free energy estimates are central to the reward formulation described in Section 3.

### 2.2. Robotics Planning and Reinforcement Learning

#### Robotics Planning Algorithms

By having estimates of free energy, we can set up various pathfinding algorithms to optimize paths on the free energy landscape. Sample-based planning methods have shown to be successful in systematically finding optimal trajectories. A gold-standard planning algorithm which many variants have been built upon is the RRT^*^ algorithm [15]. RRT^*^ uniformly samples an environment and constructs branching paths in such a way that there is guaranteed optimality with sufficient sampling. We demonstrate that our SMD sampling framework works for RRT *. Although there are other variants of RRT^*^ such as Q-RRT^*^ [14] and P-RRT^*^ [22], we use the default RRT^*^ here and the extension to any variants should be straight forward.

#### Reinforcement Learning

Because free energy is is directly connected to probability theory, we can draw several parallels between how reinforcement learning and biology both naturally optimize an objective. There have been methods that utilize reinforcement learning aspects to improve existing statistical mechanical tools. One notable example is the FAST by Zimmerman et al [2] which uses RL to encourage exploration of unexplored states for Markov-State-Modeling (MSM). Shamsi et al [4] uses RL concepts to improve adaptive sampling along a set of collective variables. Both of these methods explore the phase space with RL, but we are interested in sampling paths between two known states.

Proximal Policy Optimization (PPO) for Policy Gradient algorithms [25] is a standard deep reinforcement learning algorithm which uses a novel objective function that clips the objective function such that the new policy *π_θ_* after the gradient step is sufficiently close to the old policy 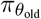. For brevity, the authors of PPO define 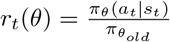and *r*(*θ*_*old*_) = 1, where *a*_*t*_ is the action, *s*_*t*_ is the state at time *t*, and *A*_*t*_ is the advantage estimate

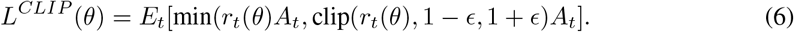

This is the objective function that we use in our methodology in order to approach monotonic improvement of the policy for stable learning on the free energy surface.

### 2.3. Machine learning for Molecular Dynamics

Modeling molecular dynamics with machine learning has been a promising direction due to developments in physics-informed models. Some works have used graph neural networks to model molecular dynamics because of the rotational and translational invariances the architecture provides [24] [16]. For generation, normalizing flows have shown to be effective for one-shot sampling of equilibrium states for molecules [1]. Conditional generative adversarial networks have also shown potential for generating low energy pathways, provided a trial set of pathways and a potential energy loss function [8]. Another relevant work uses optimal control to learn the policy of what forces to apply in order to bias the molecule to the target state [12]. Although this methodology does not use CVs, one limitation the authors mention is the transitions do not converge for molecules with naturally long transition times. We aim to learn these pathways with minimum prior knowledge of the free energy landscape except for intuition on the reduced space of CVs.

## 3 Approach

Following our outline in figure 1 and labels I-IV in this illustration, we consider a system of *N* atoms. For this system, assume we have a set of CVs that are sufficient to represent the dynamics.

**Figure 1:**
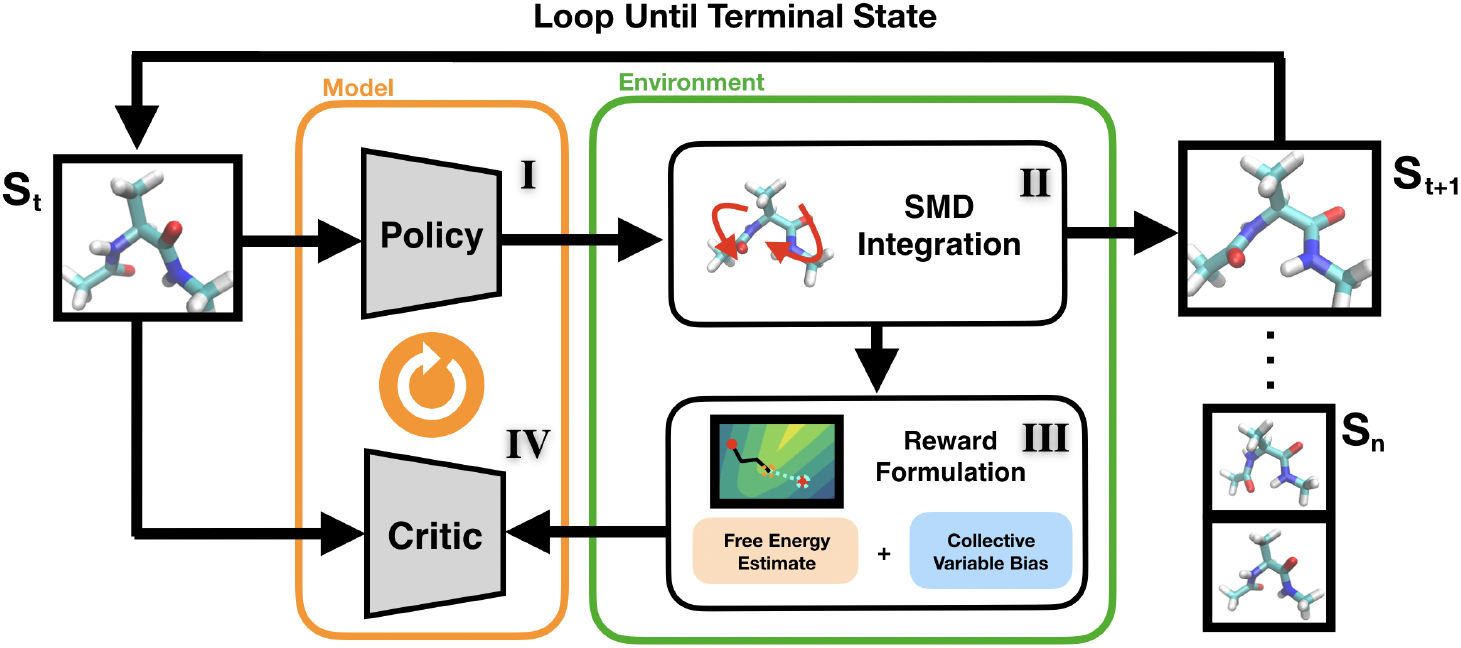
Our reinforcement learning framework uses estimates of free energy and a CV bias in order to construct a reward signal. The environment then SMD to evolve the system. We then optimize the expected returns using Proximal Policy Optimization. Parts I-IV correspond to the approach outlined in Section 3.

I. First, the SMD simulation is initialized at state A, and the policy maps the current full atom representation to a categorical distribution, where an action *a*_*t*_ can be sampled. The stochasticity of policy gradients are critical in order to encourage exploration towards alternate low energy pathways.
II. Using NAMD [5] as our environment simulator, we evolve the current state into the next state based on the action chosen by the policy. The system is evolved through a constant force stiff-spring pull along a collective variable. By simulating multiple replicas, we are able to use Jarzynski’s equality to approximate the free energy at points along the phase space. Because free energy is a state variable, we approximate the free energy difference between the starting state *A* and the current point *t* by summing the free energy differences along that path.
III. The reward *R*_*total*_ is constructed as a linear combination based on two critical parameters, *α* and *β. α* corresponds to the reward from the CV bias from state A to B. The *β* corresponds to the contribution from decreasing free energy. When *α* dominates, the model will tend towards SMD, where traveling faster to the end-state is preferred. If *β* were larger, then the trajectories would tend towards favoring convergence on intermediate low energy states.

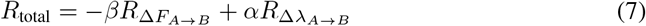 This reward function is reminiscent of the physics model equation 2, since the first term is analogous to the force field Hamiltonian *H* that relates to free energy estimates, and the second term is associated with the CV bias potential *h*_*λ*_.
IV. Based on each state-action-pair, the RL algorithm optimizes the expected long term returns at a given state. This encourages the model to tend towards states that approach the target but also least likely to run into high free energy barriers. We use the infinite horizon formulation to allow flexible timesteps and terminate the episode when it either reaches the target, or leave the bounds of the defined CVs.
V. Repeat steps 1-4 until the sampled trajectories converges to low free energy transition pathways.This process can easily be extended to that of the RRT^*^ sampling-based framework by using the free energy difference between state A and the current point as the cost.

## 4 Experiments and Results

To demonstrate that our RL reinforcement learning and RRT^*^ robotics planning algorithms can be integrated with energy estimates from SMD, we first evaluated performance on a toy system, where an analytical 2D free energy surface was known a priori. We continued to a second system, namely alanine dipeptide [23] [27], for verifying whether simultaneous reinforcement of energy and CV rewards can indeed recover pathways with correct kinetic barriers between two known end-states of a biomolecule.

### Analytical Surfaces

We prepared an artificial surface with two CVs encompassing a range of −200 to 200, commensurate to values of dihedral angles that most proteins sample. The two end states were modeled iso-energetic and separated by a barrier. A number of 8 actions were used, controlling a unit ball CV space. After training the model on 1000 iterations, we observe that the algorithm is able to learn transition pathways between two states. Depending on the strength of the contribution of each coefficient,*α* and *β*, as represented in equation 7, the pathways derived from the model will relax around high free energy barriers. We use the greatest free energy barrier traversed by the pathway as a metric to evaluate the quality of the transition pathway. Figure 2 shows how the RL algorithm converges to low energy pathways.

**Figure 2:**
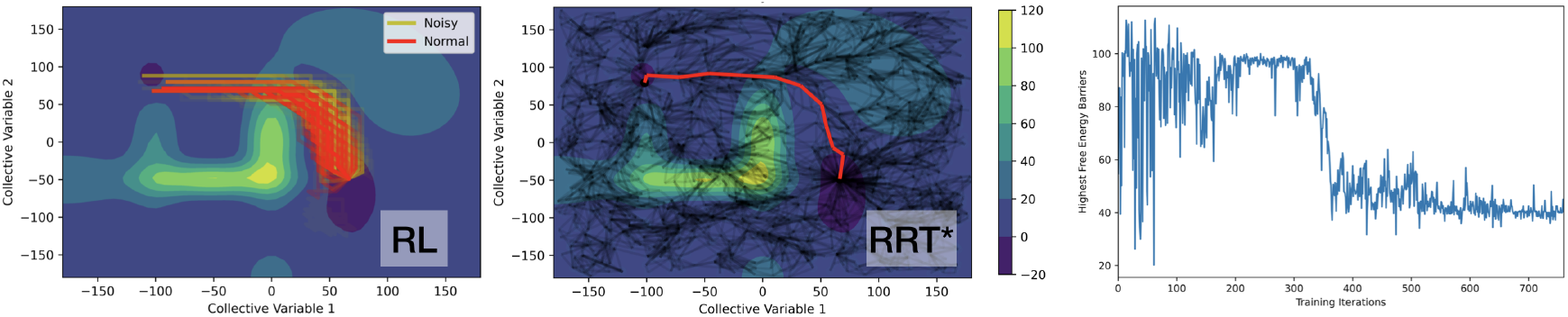
(left) RL actions with both noisy and regular forms of the Analytical Surface. (middle) RRT^*^ Algorithm. (right) Highest Free energy barriers for derived pathways plotted during training for the RL Algorithm. The coefficients used for RL here are *α* =0.4 and *β* =0.05, see equation (7).

We found that the RL algorithm not only successfully traverses the low energy barriers, but is also able to do so under significant Gaussian noise (Figure 2). This is because the RL framework works under expectation of free energy estimates. RRT^*^ is also able to recover a low energy pathway between the two known states. However, since the cost estimates for RRT^*^ do not work under expectation, we found that it did not perform as well under high amounts of noise.

### Alanine Dipeptide

We validated our model with real-world data by testing it on alanine dipeptide. We chose alanine dipeptide because the free energy surface along the backbone dihedral angles has been thoroughly understood. The RL algorithm is provided with 22×3 = 66 atomic positions, two CVs (*ϕ, ψ*) as well as two conformational states to track the movements. Figure 3 shows that both the RRT^*^ and the RL algorithms are able to recover biological transition pathways for alanine dipeptide, using 8 actions over 770 iterations. The higher barrier on the path converged to 5 kcal/mol that is in excellent agreement with several replicas of 249 ps-long free energy simulations of the system [9]

**Figure 3:**
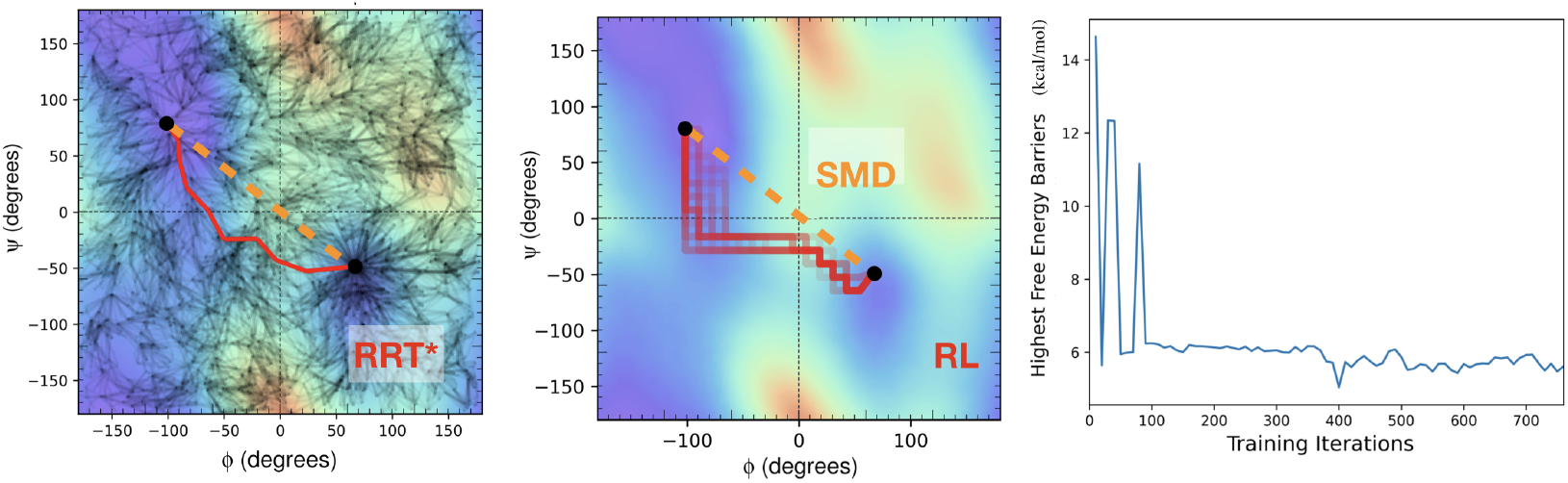
(left) RRT^*^ on the alanine dipeptide surface with direct SMD drawn for comparison (orange). (middle) and RL Algorithm for alanine dipeptide. (right) Highest free energy barriers for derived pathways during training for the RL algorithm. The coefficients used for RL here are *α* = 0.35 and *β* = 0.05.

The RRT^*^ algorithm requires significantly more sampling over the entire phase space in order to learn the minimum free energy pathway. In contrast, the RL algorithm is able to do an informed search to quickly narrow down necessary areas in phase space to traverse. However, the trajectories recovered will not be the minimum free energy pathways, but instead low free energy pathways in expectation. Similar to the analytical surface, in Figure 3 we plot the trajectories generated over CV phase space, as well as the highest free energy barriers traversed for the pathways during training.

## Discussion

Biomolecular transitions are an outcome of the mutual reinforcement between energetics and dynamics. Although the robotics planning RRT^*^ and the actor-critic PPO are different schemes, it is beneficial to demonstrate how SMD could be applied to both, showing the flexibility of the RL-SMD setup, and allowing comparison of path search algorithms. While the former performs a global search, we find that a more local search of the surface in the later is adequate to determine realistic transition paths. A strength of RL-guided SMD over brute-force pulling is that, the actions initially chosen of the policy are independent of the energy landscape. This allows the model to freely explore CV phase space and obtain samples regardless of barriers. As the rewards update, these actions are optimized to learn the landscape, avoid unphysical barriers and determine pathways.

However, there are still limitations with the current implementation. Our current algorithm has the model interface with NAMD and make gradient updates in PyTorch. This interface has a file I/O bottleneck. If we are able to train the model directly within NAMD, then the bottleneck would not be an issue. But this algorithm is also highly parallizable. Multiple SMD simulations can be deployed at once to update the main policy. Several RL algorithms do this, such as A3C [18].

## Conclusion

We demonstrate that our Jarzynski and stiff-spring motivated framework is not only able to derive low free energy transitions, but is also flexible to different kinds of path search algorithms. We believe a future direction is to use a different RL formulation of this problem. One idea is to use the entire trajectory as a given state for the policy to optimize. There are several biologically motivated metrics that can be used for assessing the entire entire pathway at once.

